# Sublethal concentrations of clothianidin affect honey bee colony behavior and interact with landscapes to affect colony growth

**DOI:** 10.1101/2020.06.05.136127

**Authors:** William G. Meikle, John J. Adamczyk, Milagra Weiss, Janie Ross, Chris Werle, Eli Beren

**Affiliations:** Carl Hayden Bee Research Center, USDA-ARS, 2000 E. Allen Rd, Tucson, AZ 85719 USA; Southern Horticultural Laboratory, USDA-ARS, P. O. Box 287, Poplarville MS 39470 USA

**Keywords:** continuous hive weight, continuous hive temperature, hive CO_2_ concentration, newly-emerged bees, neonicotinoid, pesticide residues

## Abstract

Honey bee colonies were exposed to sublethal concentrations (5 and 20 ppb) of clothianidin in sugar syrup, while control colonies were fed syrup with no pesticide. In addition to standard colony assessments of adult bees and brood, hive weight and internal temperature were monitored on a continuous basis at all sites. Experiments were conducted twice in Arizona, in successive years at the same site, and once in Mississippi, to examine the concomitant effects of weather and landscape. Adult bee masses at the Arizona site were significantly affected by clothianidin concentration. Newly-emerged bee dry weights, measured only at the Arizona site, were significantly lower for colonies fed 5 ppb clothianidin compared to the other groups. CO_2_ concentration, also only measured at the Arizona site, was higher in colonies fed 20 ppb clothianidin. Neither daily hive weight change nor colony thermoregulation were affected by clothianidin exposure. The Mississippi site had higher rainfall, more diverse land use, and a different temperature regime, and bee colonies there did not show any effects of clothianidin. These results suggest that bee colonies in more stressful environments, such as the Sonoran desert of southern Arizona, are affected more by clothianidin exposure than colonies at sites with higher rainfall and more forage. Clothianidin was also found to be, like imidacloprid, highly stable in honey in the hive environment at least over several months. These results also showed that CO_2_ concentration within the hive is potentially valuable in measuring the effects of stressors on bee health.

## Introduction

The exposure of honey bees to neonicotinoid pesticides is cosmopolitan [1]. Among neonicotinoid pesticides, thiomethoxam and its metabolite, clothianidin, are among the most popular and among the most dangerous for honey bees [2]. Clothianidin exposure has been found to affect grooming, hygienic behavior and neural gene expression [3-5]; memory processing [6]; drone semen quality [7]; and has been associated with increased P450 gene expression [8] indicating active detoxification. Clothianidin has been found in some studies to increase worker mortality [9-12], but not all studies [13] and when combined with λ-cyhalothrin has been shown to affect adult bee weight [8]. Exposure of honey bees to neonicotinoid pesticides along with other stressors, such as poor nutrition [14] or viruses [15] have been found particularly deleterious. Honey bees exposed to neonicotinoids have been found to have higher Varroa and Nosema densities [16-19] and reduced social immunity [11]. Imidacloprid, perhaps the most popular neonicotinoid, has been shown to affect brood production, queen replacement, foraging activity and winter survivorship when applied at sublethal concentrations in pollen diet [16]. When applied at very low (5 ppb) concentrations in sugar syrup, imidacloprid has been found to affect colony thermoregulation, foraging activity and adult bee maturation [20-22].

Sublethal pesticide exposure may affect aspects of honey bee ecology and social organization, but in the case of clothianidin, observations of negative impacts in managed manipulative field studies have not been consistent. Different workers have reported no effects of field-realistic concentrations of clothianidin on colony-level growth or behavior [23, 24], or on colony winter survival [25]. A large-scale study in Germany in which bee colonies were allowed to forage on oilseed crops treated with clothianidin found no effects on development of colony strength, brood success, honey yield or levels of pathogen infection [26]. Similarly, a field study involving “mini-colonies” challenged with both Nosema and clothianidin found no effect of clothianidin treatment on mortality or flight activity, and while the lifespan of *Nosema* infected bees were reduced compared to non-infected bees a combination of pesticide and pathogen did not reveal any synergistic effect [27]. Similarly, experiments with imidacloprid have also had mixed results with respect to colony growth and thermoregulation [20, 21, 28].

In this study three field experiments were conducted: two experiments at the same site in Arizona in successive years, and a third experiment at another site in Mississippi. Colony growth and activity were observed using several discrete and continuous measures in all the experiments. Additional variables were measured at the Arizona sites. The measures included those of interest to commercial beekeepers, such as the size of the adult bee and brood populations, as well as measures such as continuous hive weight and internal hive temperature that have successful detected effects on bee colonies treated with sublethal concentrations of other compounds [22, 29].

## Materials and methods

Two studies were conducted at the Santa Rita Experimental Range (SRER) south of Tucson, AZ (31°46’39”N, 110°51’46”W). The first study ran from May 2017 to March 2018 (hereafter SRER 2017-18) and the second study, from May 2018 to February 2019 (hereafter the SRER 2018-19). An additional study was conducted in Poplarville, MS (30°50’2.59”N, 89°32’52.45”W) from May 2018 to March 2019 (hereafter POPL 2018-19). An overview of the response variables is provided (Table 1).

**Table 1.** Overview of experimental design and response variables. NEB = Newly Emerged Bee.

| Experiment | No. | No. colony | NEB | Hive | Hive | Hive | Pesticide | Varroa |
| --- | --- | --- | --- | --- | --- | --- | --- | --- |
|  | colonies | assessments | mass | weight | temp. | [CO<sub>2</sub>] | residues | levels |
| SRER 2017-18 | 16 | 6 | yes | Yes | Yes | no | yes | yes |
| SRER 2018-19 | 18 | 5 | yes | Yes | Yes | yes | yes | yes |
| POPL 2018-19 | 15 | 3 | no | Yes | Yes | no | no | yes |

### Syrup preparation

Control (0 ppb clothianidin) sucrose solution was mixed at 1:1 w:w (e.g. 500 g sucrose:500mL distilled water). Sucrose was added to distilled water in a 5-gallon bucket and mixed using an electric drill with a mortar mixing attachment until sugar was completely dissolved. Sucrose solution for solutions with clothianidin (PESTANAL, CAS# 210880-92-5) was mixed in the same manner but 50mL was withheld (thus “short”) to allow for the added volume of respective clothianidin spikes. 500 g of sugar is dissolved in 450 mL of distilled water to allow for the addition of a 50 mL spike to achieve 1 kg of treatment solution. 950 g of “short” sugar solution was transferred to a Nalgene bottle, then the spike added to each individual bottle. A 10 ppm clothianidin stock solution was made by dissolving 1.0 mg of clothianidin, in 100 mL of distilled water, using a mixing bar but without heat. To avoid problems with static electricity, the clothianidin was weighed into a small, nonreactive plastic receptacles and those receptacles were placed in the solution, the solution stirred, and the receptacles removed after confirming the clothianidin had dissolved. For the 5 ppb solution: 0.5 mL of the stock solution was mixed into 49.5 mL of distilled water to achieve 50 mL of spike solution, which was then added to 950g of the short sucrose solution to achieve 1 kg of 5 ppb clothianidin syrup. For the 20 ppb solution (only in 2nd experiment) 2.0 mL of stock solution was mixed into 48.0 mL of distilled water, and that solution added to 950 g of the short solution to achieve 1 kg of 20 ppb clothianidin syrup.

### SRER 2017-18 experiment

On 20 April, 2017, 24 bee colonies were established from packages (C.F. Koehnen & Sons, Inc., Glenn, CA 95943) of approximately 1 kg honey bees in painted, 10-frame, wooden Langstroth boxes (43.7 l capacity) (Mann Lake Ltd,) with migratory wooden lids. At establishment each colony was given 4 full or partial frames of capped honey, 2 frames of drawn but empty comb, 2 frames of partially drawn with some capped honey, 3 frames of foundation and a 1-frame feeder. Hives were placed on stainless steel electronic scales (Tekfa model B-2418 and Avery Weigh-Tronix model BSAO1824-200) (max. capacity: 100 kg, precision: ±20g; operating temperature: −30°C to 70°C) and linked to 16-bit dataloggers (Hobo UX120-006M External Channel datalogger, Onset Computer Corporation, Bourne, MA, USA) with weight recorded every 5 minutes. The scales were powered by deep-cycle batteries connected to solar panels. The system had an overall precision of approximately ±20 g. Hives were arranged in a circular pattern around a central box that contained the batteries and electronic gear. Hives within such a group were 0.5-1 m apart and groups were >3 m apart.

Colonies were all fed 2 kg sugar syrup (1:1 w:w) and 250 g pollen patty, made at a ratio of 1: 1: 1 corbicular pollen (Great Lakes Bee Co.): granulated sugar: drivert sugar (Domino Foods). The apiary was surrounded by native, unmanaged plants, including mesquite (*Prosopis* spp.), creosote (*Larrea* spp.), cactus (mainly *Opuntia* spp.) and wildflowers. No commercial agriculture exists within a 10 km of the apiary. On 10 July a temperature sensor (iButton Thermochron, precision ±0.06°C) enclosed in plastic tissue embedding cassettes (Thermo Fisher Scientific, Waltham, MA) was stapled to the center of the top bar on the 5^th^ frame in the bottom box of each hive and set to record every 15 min. The same day, pieces of slick paperboard coated with petroleum jelly and covered with mesh screens were inserted onto the hive floor to monitor *Varroa* mite fall within the hive. The boards were removed 2 days later, and the number of mites counted on each board. Infestation levels of *Varroa* were again monitored during and post treatment. Colonies were treated with amitraz (Apivar) on 19 October.

Hives were assessed on 12 July, and approximately every 5-6 weeks thereafter until November, using a published protocol (see [22, 29, 30]). Briefly, the hive was opened after the application of smoke, and each frame was lifted out sequentially, gently shaken to dislodge adult bees, photographed using a 16.3 megapixel digital camera (Canon Rebel SL1, Canon USA, Inc., Melville, NY), weighed on a portable scale (model EC15, OHaus), and replaced in the hive. Frame photographs were analyzed later in the laboratory (see below). During this first assessment (but not subsequent assessments), all hive components (i.e. lid, inner cover, box, bottom board, frames, entrance reducer, internal feeder) were also shaken free of bees and weighed to yield an initial mass of all hive components. At the initial inspection, 3-5 g of wax were collected from each hive into 50 ml centrifuge tubes and stored at −80°C; samples collected in September, prior to treatment, were pooled and subjected to a full panel analysis for residues of pesticides and fungicides, from all major classes, by the Laboratory Approval and Testing Division, Agricultural Marketing Service, USDA. Samples from later assessments were pooled within treatment group and subjected only to neonicotinoid residue analysis. Hives were assessed again 13 February 2020 and finally on 29 March.

Newly-emerged bees (NEBs) were sampled by pressing an 8 cm x 8 cm x 2 cm cage of wire mesh into a section of capped brood, then returning the following day to collect NEBs that had emerged within the cage over the previous 24 h. The NEBs were then placed in a 50 mL centrifuge tube, frozen on dry ice, and stored at −80°C. At the laboratory, 5 bees per hive per assessment date were placed in Eppendorf tubes, weighed, dried for 72 h at 60°C, then re-weighed to determine average wet and dry weight per bee. NEBs were collected on 12 July and 24 August 2017 (brood levels were too low in October 2017 for sampling).

After the first assessment, hives were ranked in terms of adult bee mass and then randomly assigned to treatment group, ensuring that the average bee masses per group were approximately equal and after eliminating assignments with excessive clumping by treatment. Just prior to treatment all broodless frames containing honey and/or pollen were replaced with frames of empty drawn comb collected earlier from the same apiary. Colonies were then fed 3 kg syrup twice per week from 14 July to 21 August, with clothianidin concentrations depending on their treatment group. Hives were assessed approximately every 5-6 weeks thereafter until November, and again in February and March. Hives were inspected from lowest to highest concentration treatments, and all equipment cleaned or changed between treatment groups.

### SRER 2018-19 experiment

The 2017-18 experiment was repeated. On 11 April, 2018, 24 bee colonies were established from packages from the same supplier again into Langstroth boxes at the same location, with approximately the same initial assortment of drawn comb and food resources. Hives were again placed on the hive scales, powered and monitored in the same manner. Colonies were fed in the same manner as before. Temperature sensors were installed on 28 June. Varroa mite fall onto adhesive boards was monitored 6-9 July. Hives were assessed on 5 July in the same manner as before, and wax, honey, and NEBs were sampled. NEBS were sampled on 6 July, 23 August and 4 October, 2018. CO_2_ probes (Vaisala), calibrated for 0-20% concentrations, were installed in five hives in each treatment group and set to record CO_2_ concentration every 5 minutes.

After the first assessment, hives were ranked in terms of adult bee mass and assigned to treatment groups in the same manner as the previous year. Again, just prior to treatment all broodless frames containing honey and/or pollen were replaced with frames of empty drawn comb collected earlier from the same apiary. Colonies were fed 3 kg sugar syrup twice per week from 12 July to 20 August with the same pesticide concentrations as the previous year.

Varroa infestation levels were again monitored at the end of August and again at the beginning of November. Colonies were treated with amitraz (Apivar®) on 19 October. Hives were assessed approximately every 5-6 weeks thereafter until November, and then in February, at which point the experiment was ended. Hives were inspected from lowest to highest concentration treatments, and all equipment cleaned or changed between treatment groups.

### POPL 2018-19 experiment

Full bee colonies, each comprised of two “deep” boxes as described above, were obtained from a local bee supplier (Gunter Apiaries, Lumberton MS) as nucleus colonies the previous year. Colonies were placed on hive scales (Tekfa model B-2418) on 16 May 2018. Colonies were assessed, using the methods described above, on 11 July 2018 and temperature sensors (iButtons) were installed on 12 July 2018. Frames of honey were removed on 18 July and colonies were randomly placed in treatment groups. Treatment feeding commenced 24 July, lasting 31 August, using the same concentrations and amounts as described above. Colonies were not fed pollen patty because sufficient pollen was available. Colonies were assessed again 20 September 2018 and finally on 27 March 2019. Samples of 300 bees were collected on 7 May 2018, washed in 70% ethanol and the Varroa mites counted. Colonies were treated for Varroa (Checkmite, Mann Lake Ltd) on 28 June 2018. The apiary site was assessed using the National Agricultural Statistical Service (NASS) Cropscape web site (https://nassgeodata.gmu.edu/CropScape) to obtain acreage estimates for all land use categories within an approximately 1.8 km radius of the apiary.

### Data analysis

The area of sealed brood per frame was estimated from the photographs using ImageJ version 1.47 software (W. Rasband, National Institutes of Health, USA) or CombCount [31]; this method has been described in other publications (e.g. [29, 30, 32]).

The total weight of the adult bee population was calculated by subtracting the combined weights of hive components (i.e. lid, inner cover, box, bottom board, frames, entrance reducer, internal feeder) obtained at the start of the experiment (model EC15, OHaus) from the total hive weight recorded the midnight prior to the inspection.

Honey bee colony survivorship was analyzed using Proc LifeReg (SAS Inc. 2002). Survivorship curves were generated for each treatment group within each experiment. Treatments compared using ANOVA (α=0.05) (Proc Glimmix, SAS Inc. 2002) with respect to three parameters: 1) the 30^th^ percentile; 2) the 50^th^ percentile; and 3) a shape variable calculated by subtracting the 40^th^ from the 30^th^ percentile. Daily hive weight change was calculated as the weight change from midnight of a given day to 23 h 55 min later. Continuous temperature data were divided into daily average data and within-day detrended data. Detrended data were obtained as the difference between the 25 hour running average and the raw data. Sine curves were fit to 3-day subsamples of the detrended data, taken sequentially by day (see [32]). Curve amplitudes, representing estimates of daily variability, were reduced to a data point every 3 days, to ensure no overlap between data subsamples, for repeated measures MANOVA analyses. CO_2_ concentration data were treated in the same fashion.

Adult bees, brood surface area, daily hive weight change, internal hive temperature average and variability (i.e. amplitudes of fit sine curves) and CO_2_ concentration average and variability were used as response variables in repeated-measures MANOVA (Proc Glimmix, SAS Inc. 2002) with Treatment, Sampling date, Experiment and Day, and all 2-way interactions, as fixed effects and with pre-treatment values as a covariate to control for pre-existing differences. Analyses of hive weight and temperature were limited to approximately 3 months after the end of treatment to focus on the active season, and initially consisted of omnibus tests that included all three field experiments followed by analyses within each experiment. The reason for this is that effects that are significant in one trial might not be so in another, or might be significant but in a contrary fashion. CO_2_ concentration data were only collected in the SRER 2018-19 experiment.

NEB data were analyzed with Treatment, Sampling date and their interaction, with the July values as a covariate. Varroa fall were analyzed within each SRER experiment, with the pre-treatment values used as covariates where applicable. Varroa alcohol wash data for POPL 2018-19 were analyzed separately.

Ambient temperature, rainfall and ambient CO_2_ data were obtained for Arizona: AmeriFlux US-SRM Santa Rita Mesquite, doi:10.17190/AMF/1246104; and temperature and rainfall data for Mississippi: National Environmental Satellite, Data, and Information Service, National Oceanic and Atmospheric Administration, Poplarville Experimental Station, MS US USC00227128.

## Results

### Hive survivorship

No significant differences were observed among treatment groups with respect to hive survivorship for any of the experiments (p=0.40 for the 30^th^ percentile, p=0.34 for the 50^th^ percentile, and p=0.32 for the difference between the 30^th^ and 40^th^ percentiles) (Figure 1). Four of 6 colonies died in the clothianidin 20 ppb treatment group during the course of the SRER 17-18 experiment, compared to 2 in the Control treatment group and 1 in the clothianidin 5 ppb treatment group, while in SRER 2018-19 both the clothianidin 20 ppb and Control groups lost 2 colonies while the clothianidin 5 ppb group lost 3. POPL 2018-19, the treatment groups with clothianidin both lost 2 of 5 colonies while the Control group lost 3.

**Fig. 1.**
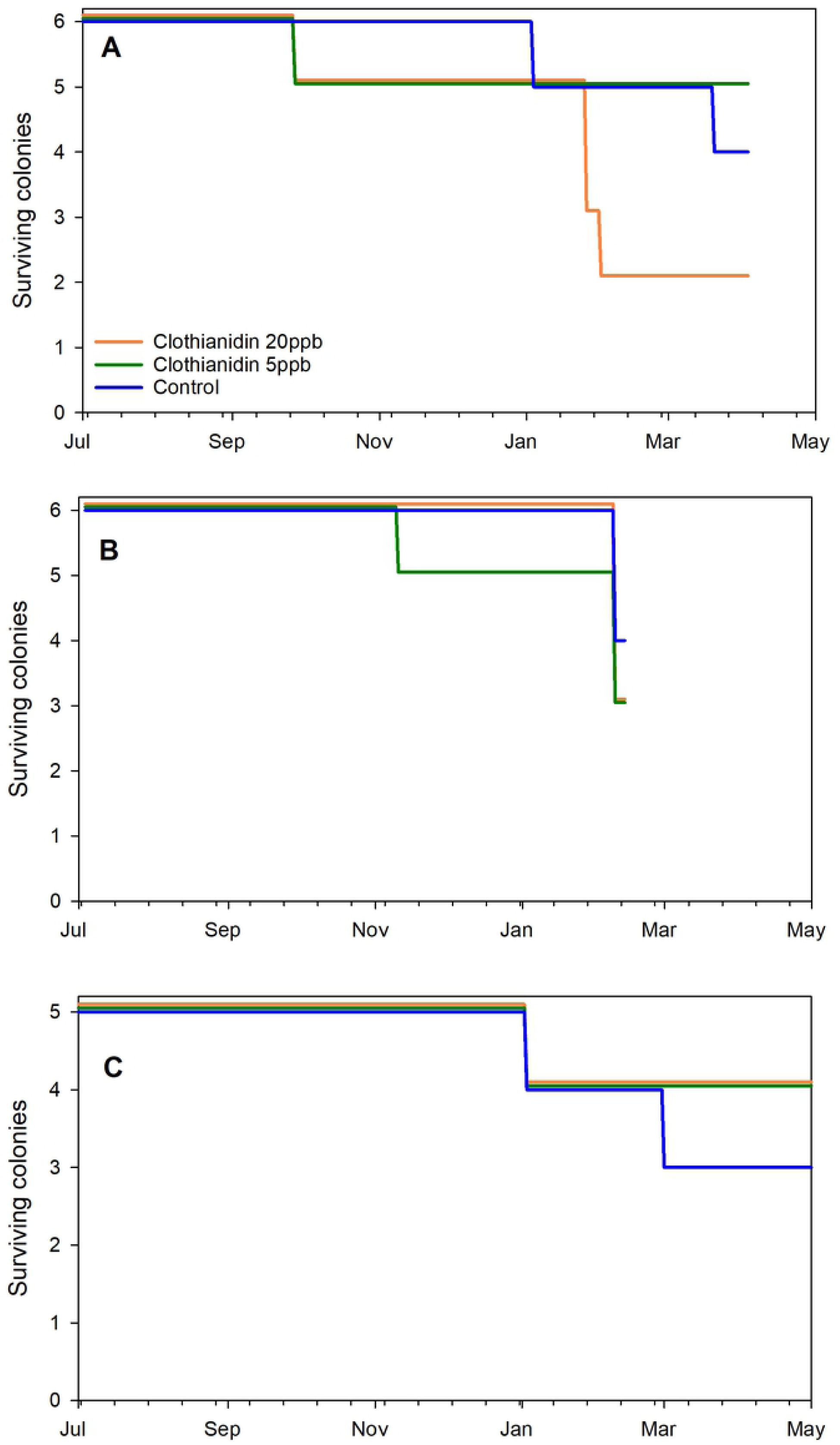
Honey bee colony survivorship for each of 3 treatment groups. A) SRER 2017-18; B) SRER 2018-19; C) POPL 2018-19.

### Adult bees

Hive evaluations were conducted on different schedules between the two experimental sites (SRER and Poplarville), so analyses for the two sites were conducted separately. Treatment effects were significant at the SRER site (p=0.0456) (S1 and S2 Tables); pairwise contrasts did not reveal any significant differences at the p=0.05 level among treatment groups (p=0.0571 for the contrast between clothianidin 20 ppb and control groups) (Figure 2). Treatment effects were not significant in the Poplarville experiment (p=0.62).

**Fig. 2.**
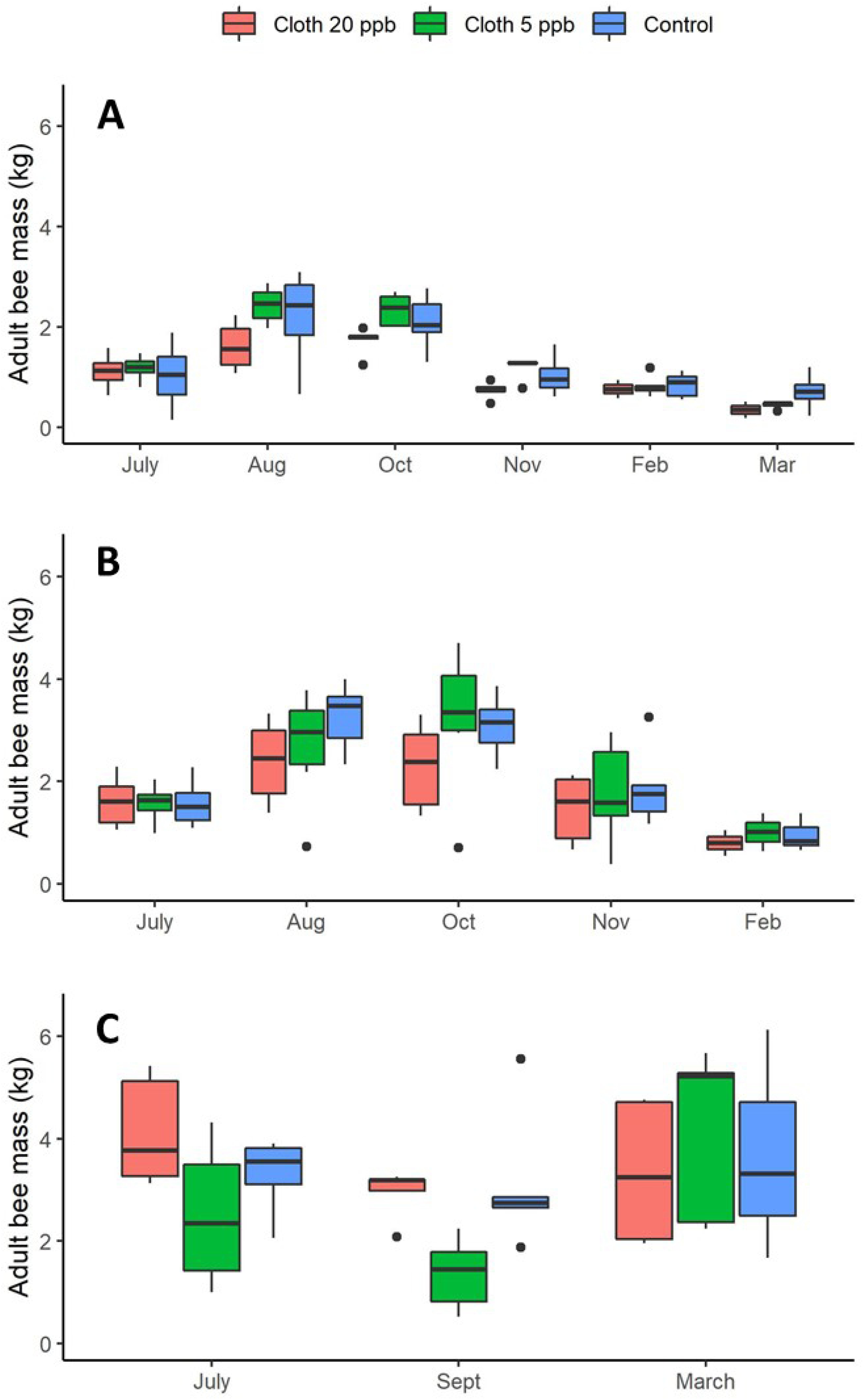
Average adult bee mass (kg) per colony for each of 3 treatment groups: Clothianidin 20 ppb (blue), clothianidin 5 ppb (orange), control (gray). A) SRER 2017-18; B) SRER 2018-19; C) POPL 2018-19. Boxes are defined as 1.58 * IQR / n^0.5^, where IQR is the inter-quartile range and n is the number of data points.

### Brood

Considering the brood surface area by site, as described above, treatment had no significant effect at either the SRER site (p=0.55) or the Poplarville site (p=0.38) (Figure 3, S3 Table).

**Fig. 3.**
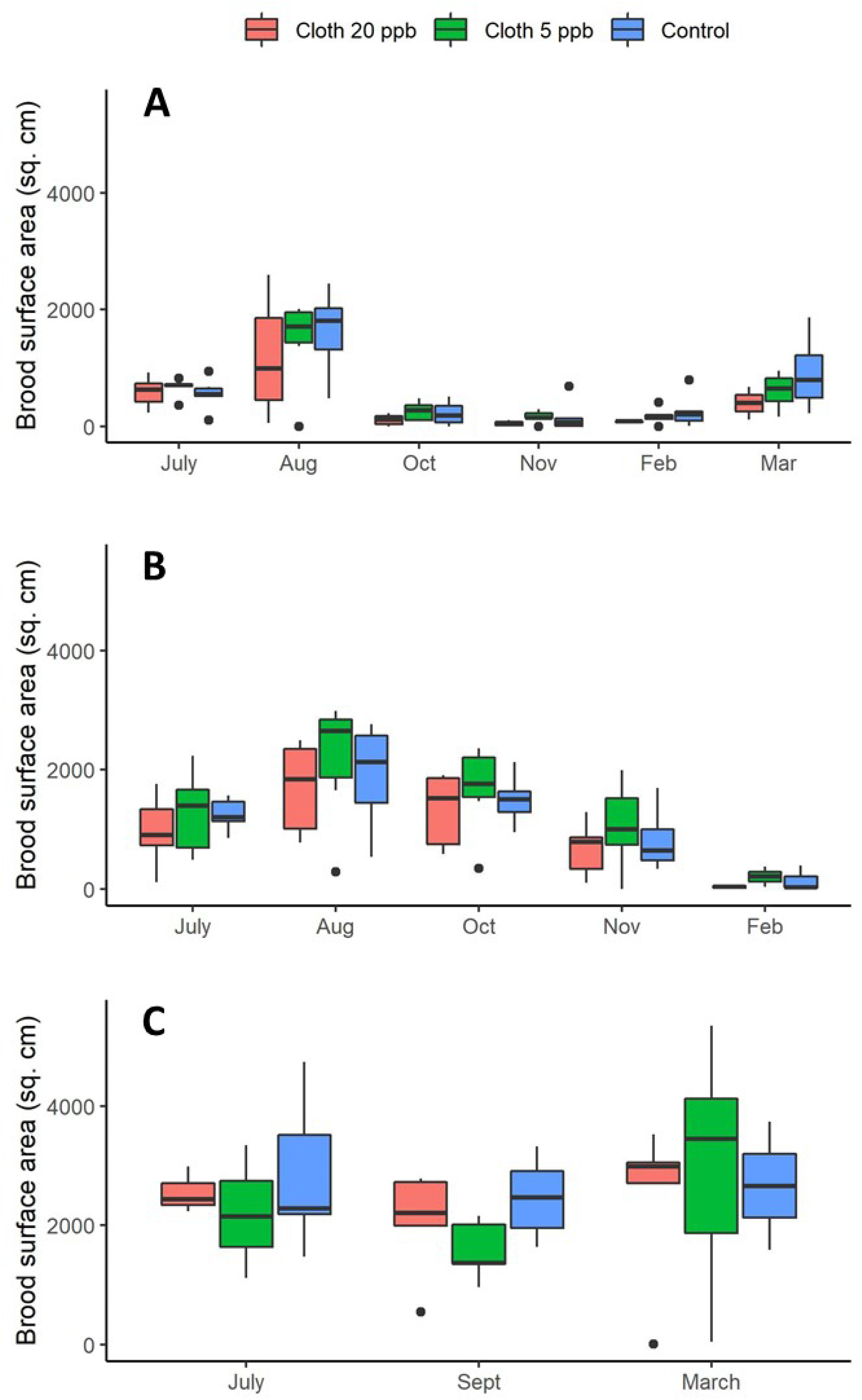
Average brood surface area (cm^2^) per colony for each of 3 treatment groups: Clothianidin 20 ppb (blue), clothianidin 5 ppb (orange), control (gray). A) SRER 2017-18; B) SRER 2018-19; C) POPL 2018-19. Boxes are defined as 1.58 * IQR / n^0.5^, where IQR is the inter-quartile range and n is the number of data points.

### Newly Emerged Bees

Data for the SRER 2017-18 experiment were only available for July (pre-treatment) and August (immediately post-treatment) and treatment was not significant (p=0.19) (S4-S7 Tables). With respect to the 2018-19 experiment, treatment had a significant effect (p=0.0046) on NEB dry weight for the August and October samples collected in 2018 (Figure 4). Neither sampling date nor the interaction term were significant, indicating the relationships remained largely the same between the two sampling dates. Pairwise contrasts showed that average NEB dry weight from the clothianidin 5 ppb treatment group were significantly smaller than those from the control group (p=0.0054) and the clothianidin 20 ppb group (p=0.0310). The control and clothianidin 20 ppb groups were not significantly different.

**Fig. 4.**
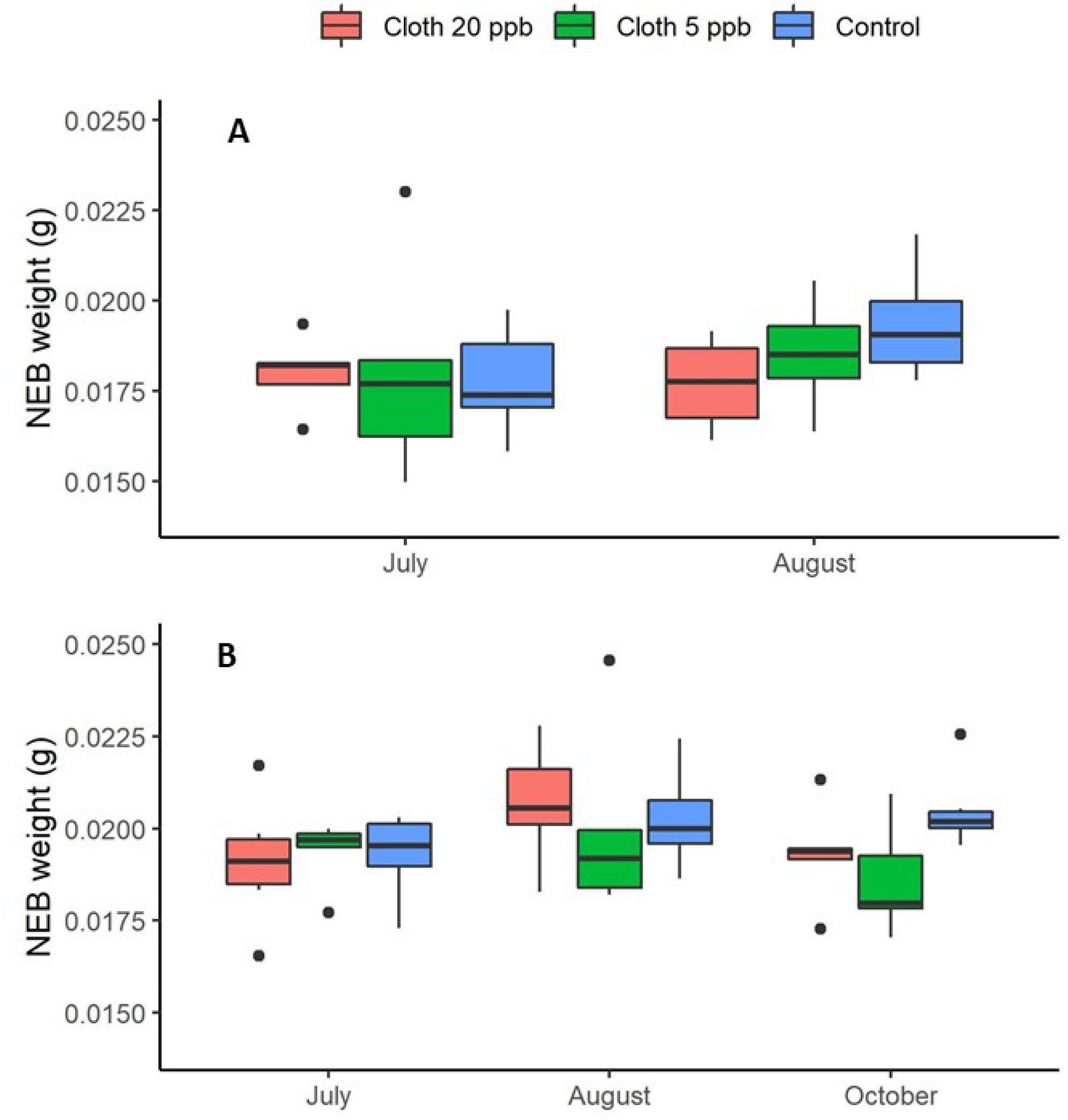
Average Newly Emerged Bee (NEB) dry weights for each of 3 treatment groups: Clothianidin 20 ppb (blue), clothianidin 5 ppb (orange), control (gray). A) SRER 2017-18 (2 sampling dates); B) 2018-19 experiment (3 sampling dates). Boxes are defined as 1.58 * IQR / n^0.5^, where IQR is the inter-quartile range and n is the number of data points.

Considering the SRER 2017-18 and 2018-19 experiments together for August (the only sampling date that both experiments had in common), and again using pre-treatment values as a covariate, treatment was significant (p=0.0413), while year and the treatment x year interaction were not. Pairwise contrasts showed that the control NEB dry weights were significantly larger than those for clothianidin 5 ppb (p=0.0440).

### Daily hive weight change

Because daily hive weight change was monitored in the same manner among all sites and years, all experiments could be included in the same analyses. Considering all experiments together during the first 60 days after the end of the treatment period, no significant treatment effects were observed (p=0.57) (Figure 5) (S8-S13 Tables). However, experiments were different from each other (p<0.0001) and hives in the SRER 2017-18 experiment had significantly lower daily weight gain than hives in either the SRER 2018-19 or POPL 2018-19 experiments (p<0.0001 and p=0.0205, respectively). When experiments were considered separately, treatment effects were significant for the SRER 2018-19 study (p=0.0256) and the clothianidin 5 ppb treatment group gained more weight per day on average than the clothianidin 20 ppb treatment group (p=0.0254). Treatment effects were not significant for the SRER 2017-18 (p=0.46) or the Poplarville 2018-19 study (p=0.11).

**Fig. 5.**
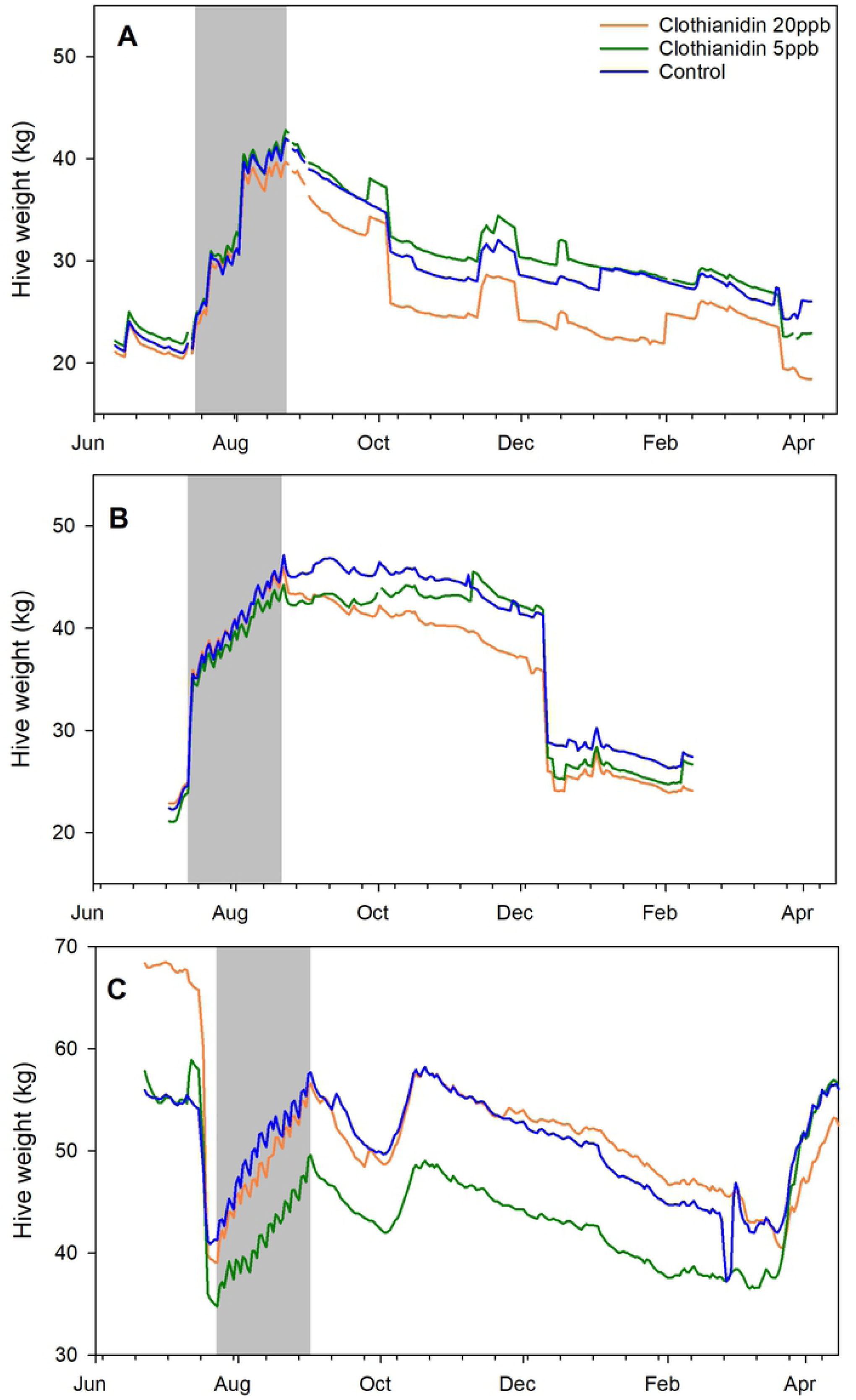
Average colony weight (kg) per hour for each of 3 treatment groups: Clothianidin 20 ppb (blue), clothianidin 5 ppb (orange), control (gray). A) SRER 2017-18; B) SRER 2018-19; C) POPL 2018-19.

### Hive temperature

Internal hive temperature and temperature amplitude from the end of treatment in August until the end of the following December, to capture the end of the annual active season, were used as response variables. No significant differences in average internal hive temperature (Figure 6) or temperature variability (Figure 7) were observed with respect to treatment, either in an omnibus analysis including all three field trials or in analyses considering each experiment separately (S14-S23 Tables). Average temperature was significantly different among experiments (p=0.0367), and in pairwise contrasts only temperatures between SRER 2017-18 and SRER 2018-19 experiments were significantly different (p=0.0362). Temperature variability was likewise different among experiments (p<0.0001) and pairwise contrasts indicated significant differences in all pairwise contrasts of experiments (all p < or =0.0001).

**Fig. 6.**
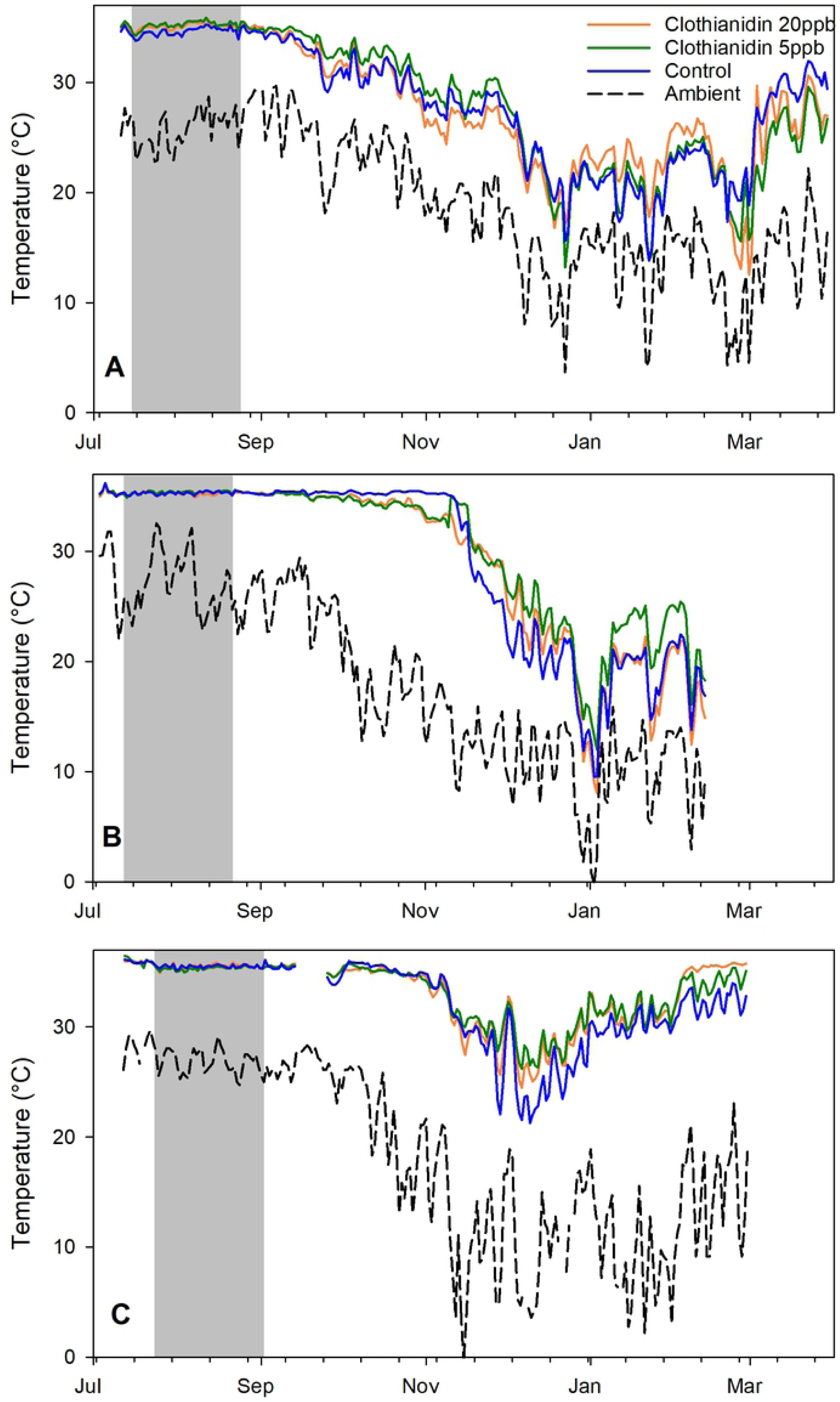
25 hour running average internal hive temperature (°C) per hour for each of 3 treatment groups compared to ambient temperatures. A) SRER 2017-18; B) SRER 2018-19; C) POPL 2018-19.

**Fig. 7.**
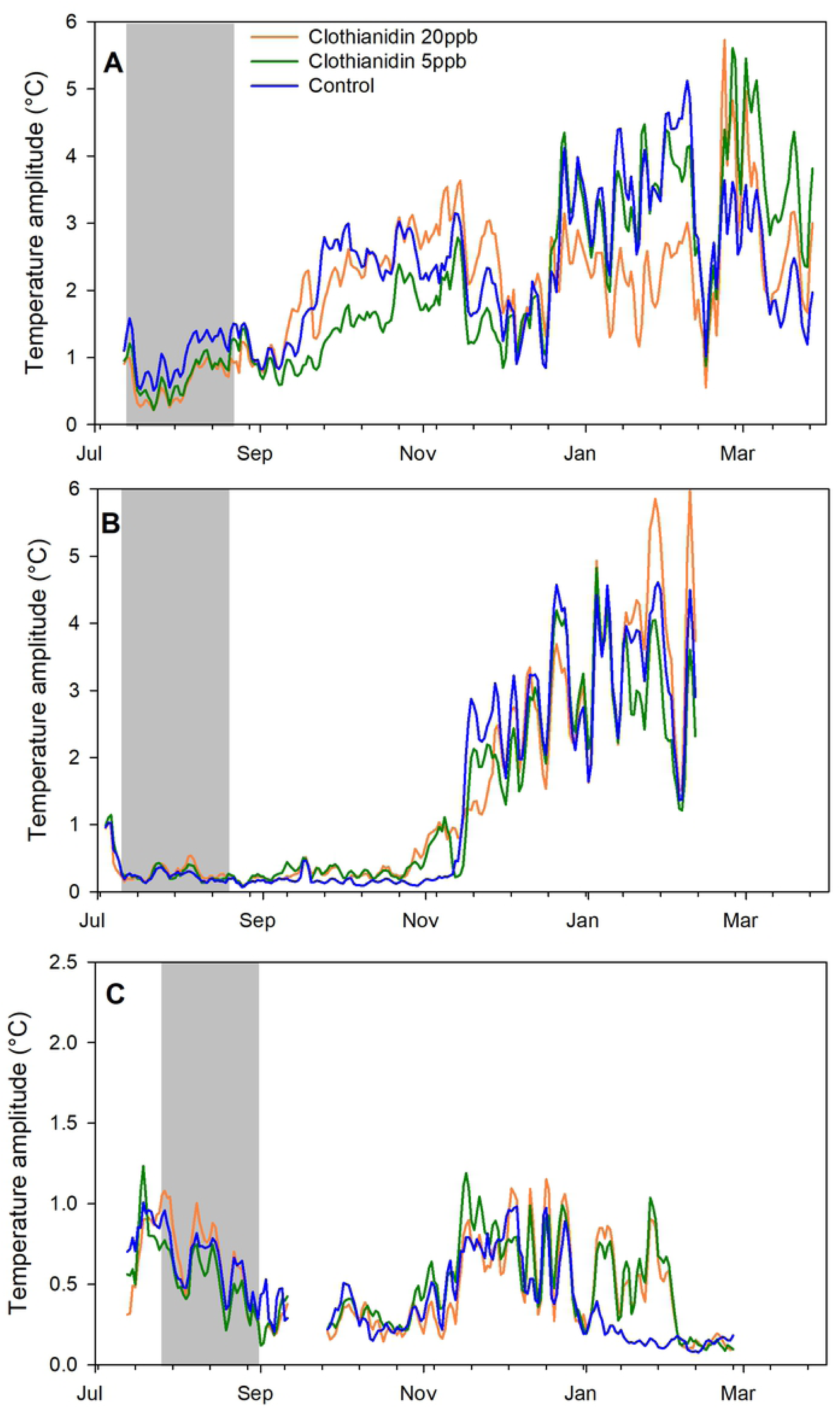
Average daily amplitudes of sine curves fit to within-day temperature changes per day (see text for details) for each of 3 treatment groups. A) SRER 2017-18; B) SRER 2018-19; C) POPL 2018-19.

### CO_2_ concentration

Treatment had a significant effect (p=0.0073) on average CO_2_ concentrations within the hive for at least the first two months after the end of the treatment period, from 31 August to 31 October (Figure 8, S24 and S25 Tables). Pairwise contrasts indicated that hives in the clothianidin 20 ppb treatment group had significantly higher CO_2_ concentration than either the clothianidin 5 ppb group (p=0.0064) and the control group (p=0.0405). Treatment did not have a significant effect on CO_2_ concentration variability (amplitude) (p=0.13). Daily amplitudes within the hives ranged across treatment groups from 1933 to 2441 ppm, whereas amplitudes of ambient CO_2_ averaged 49 across the same time period.

**Fig. 8.**
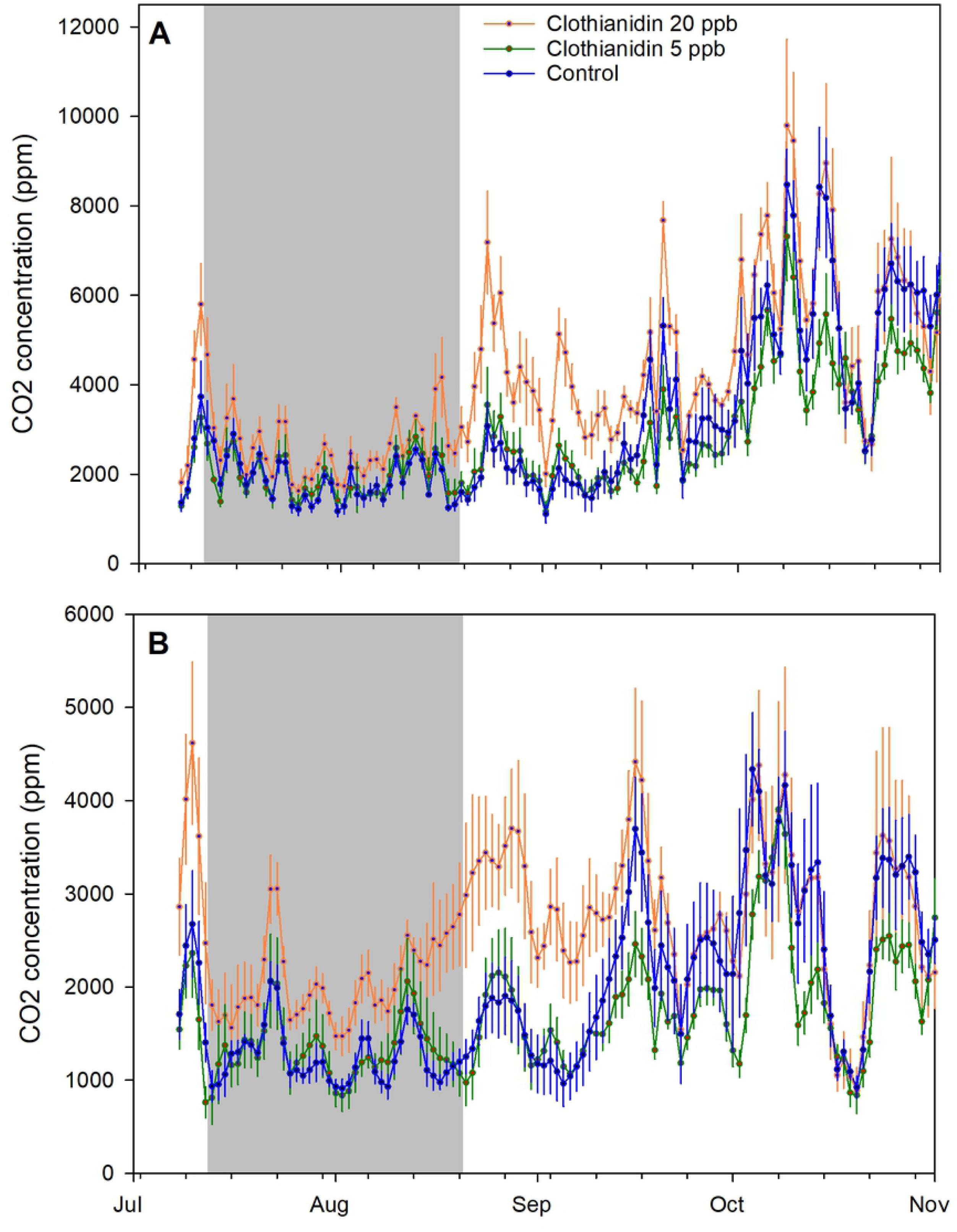
Running average CO_2_ concentrations, and daily amplitudes of sine curves fit to within-day CO_2_ concentration changes per day (see text for details), for each of 3 treatment groups for the SRER 2018-19 experiment. A) 25 hour running average; B) daily amplitudes.

### Varroa density

Varroa mite fall per hive was not affected by treatment group either in the SRER 2017-18 (p=0.48) or SRER 2018-19 (p=0.82) experiments (Table 2, S26 and S27 Tables).

**Table 2.** Mite infestations per experiment. Mite levels in the two SRER experiments were calculated as the number of mites fallen per colony per day; mite levels in the POPL 2018-19 experiment were calculated as number of mites per 100 bees from samples of 300 bees.

| Treatment | SRER 2017-18 |  | SRER 2018-19 |  |  | POPL 2018-19 |
| --- | --- | --- | --- | --- | --- | --- |
|  | July | October | July | October | November | May |
| Clothianidin 20 ppb | 2.7±0.9 | 7.8±4.2 | 3.9±1.3 | 18.7±7.1 | 17.5±6.7 | 8.9±8.1 |
| Clothianidin 5 ppb | 2.0±0.8 | 19.8±11.6 | 3.4±1.7 | 16.9±6.7 | 37.1±12.9 | 3.7±1.1 |
| Control | 1.5±0.7 | 15.3±10.0 | 6.6±2.3 | 14.3±4.1 | 38.0±18.7 | 5.6±1.9 |

### Landscape analysis

Analysis of the Poplarville landscape using CropScape yielded the following usage patterns within foraging distance of the Poplarville apiary (Table 3).

**Table 3.**
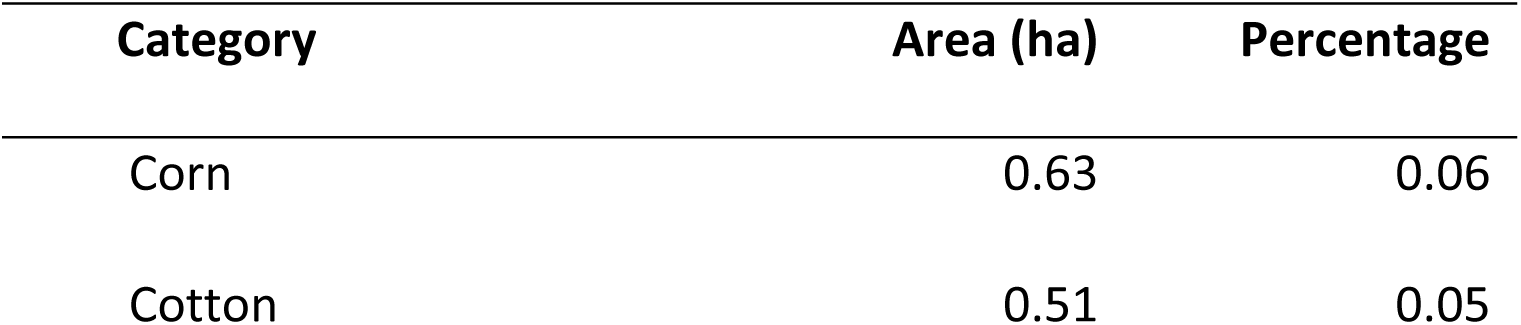
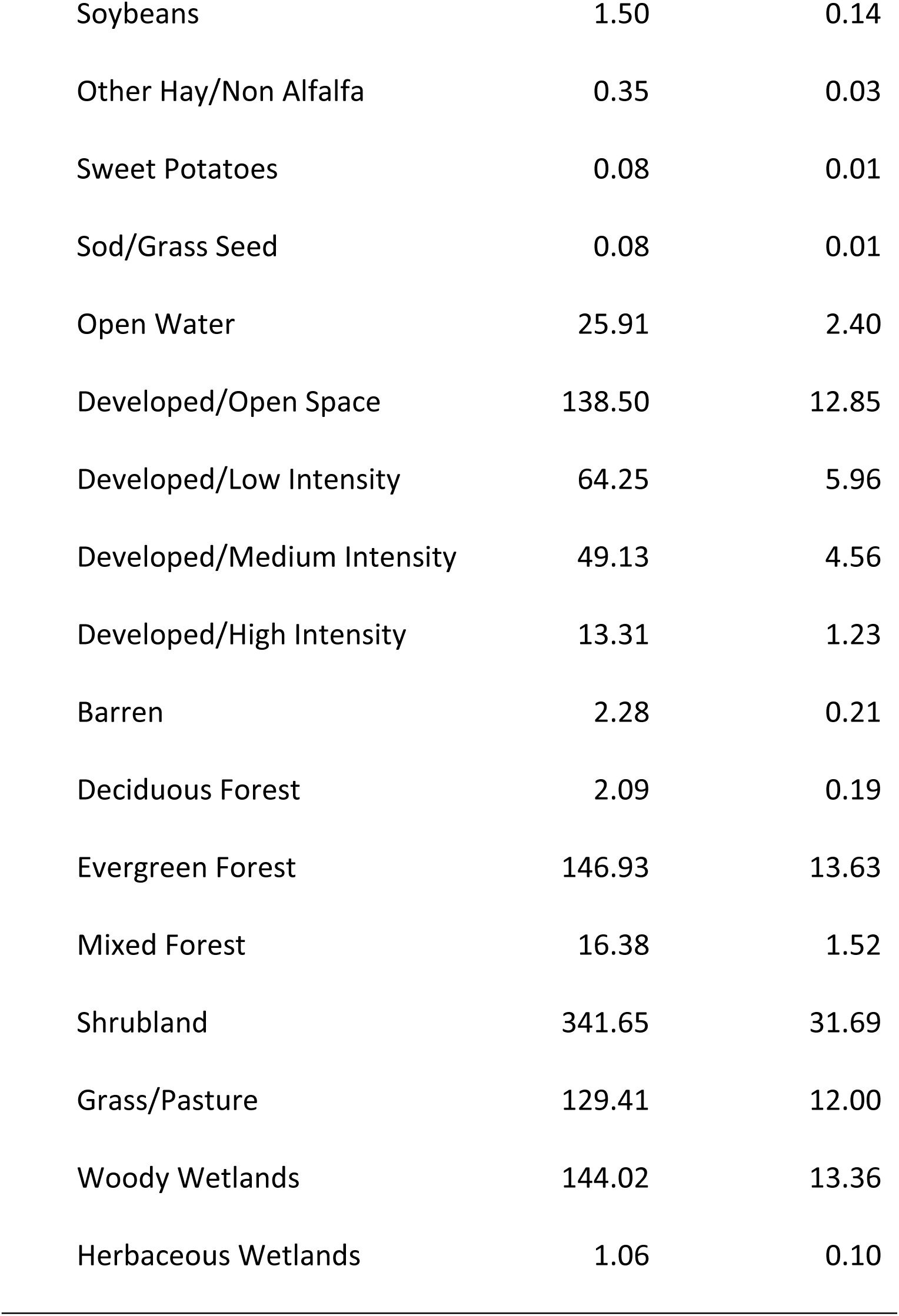
Estimated surface area and percentage area for a circle with a radius of approximately 1.8 km (= approximately 1018 ha) of land around the MS apiary in this study according to the Cropscape web site (see text for details).

### Pesticide residues

Residues in honey other than clothianidin were limited to thymol and trace amounts of 2,4-dimethylphenyl formamide (2,4-DMPF) in one sample (Table 4). Wax samples had many compounds but the residue concentrations were very low compared to acute contact LD_50_ (Table 5).

**Table 4.** Concentrations of clothianidin and thymol in honey and syrup samples across treatment groups for the two experiments in Arizona. Values are parts per billion.

| Year | Treatment group | Matrix | Thymol | Clothianidin |  |  |  |
| --- | --- | --- | --- | --- | --- | --- | --- |
|  |  |  |  | 7/10/2017 | 8/24/2017 | 11/9/2017 | 2/21/2018 |
| 2017 | Cloth_20 | Honey | 15 | 0 | 153 | 107 | 103 |
|  | Cloth_05 | Honey | 23 | 0 | 42 | 34 | 18 |
|  | Control | Honey | 89* | 0 | 0 | 0 | 0 |
|  | Cloth_20 | Syrup |  | 46 |  |  |  |
|  | Cloth_5 | Syrup |  | 12 |  |  |  |
|  |  |  |  | 7/9/2018 | 8/22/2018 |  |  |
| 2018 | Cloth_20 | Honey | 55 | 0 | 22 |  |  |
|  | Cloth_05 | Honey | - | 0 | trace |  |  |
|  | Control | Honey | - | 0 | 0 |  |  |
|  | Cloth_20 | Syrup |  | 33 |  |  |  |
|  | Cloth_5 | Syrup |  | 12 |  |  |  |

\*Trace amounts of 2,4-DMPF were also detected.

**Table 5.** Pesticide concentrations in wax samples collected pre-treatment in the 2017-18 experiment in Arizona. Values are parts per billion. “LOD” means Limit of Detection; “DMPF” is dimethylphenyl formamide. Data on acute contact LD_50_ were obtained from the Pesticide Properties Database (https://sitem.herts.ac.uk/aeru/ppdb/en/atoz.htm) and converted from µg per bee to ppb assuming an average bee mass of 0.1g.

| Compound | LOD | Contact | 2017-18 Treatment group | 2018-19 |
| --- | --- | --- | --- | --- |

|  |  | LD<sub>50</sub> | Cloth_20 | Cloth_5 | Control | Composite |
| --- | --- | --- | --- | --- | --- | --- |
| 2,4-DMPF | 1.5 | 7.50E+05 | 7 | 14 | 32 | 56 |
| Boscalid | 5 | >2.00E+06 | 5 | - | - | - |
| Carbendazim | 2 | >5.00E+05 | 24 | 27 | 37 | trace |
| Chlorthal-dimethyl | 2 | >1.00E+06 | - | trace | - | - |
| Coumaphos oxon | 1 | 5.93E+04 | 1 | 1 | 1 | trace |
| Cyprodinil | 2 | >7.84E+06 | - | trace | - | - |
| Diuron | 1 | >1.02E+06 | 3 | 1 | 2 | trace |
| Fenamidone | 1 | >2.57E+05 | trace | - | - | - |
| Fenazaquin | 1 | 1.21E+04 | - | 2 | 1 | - |
| Fenpyroximate | 3 | 1.58E+05 | 5 | - | 3 | trace |
| Flumeturon | 1 | >1.00E+06 | - | - | - | trace |
| Fluvalinate | 25 | 4.32E+04 | trace | trace | trace | trace |
| Hexythiazox | 2 | >2.00E+06 | trace | trace | trace | trace |
| Pendimethalin | 50 | 1.00E+06 | - | - | trace | - |
| Piperonyl butoxide | 6 | NA | 36 | 59 | trace | trace |
| Propargite | 2 | 4.79E+05 | 32 | 19 | 29 | 7 |
| Thymol | 2 | NA | 799 | 991 | 2190 | 1470 |
| Trifluralin | 10 | >1.00E+06 | - | - | - | trace |

### Rainfall

Monthly precipitation during each of the trials is provided (Figure 9). Total precipitation differed greatly between the Arizona site and the Mississippi site, as well as between years at the Arizona site. The Mississippi site received 1395 mm during the experimental period whereas the Arizona site received an average of 413 mm. Precipitation was clearly more constant over the year in Mississippi than in Arizona, which has strong seasonality. At the Arizona site, 286 mm of precipitation fell during the 2017-18 experiment while 540 mm fell the following year, an increase of 89%.

**Fig. 9.** Monthly rainfall during each field experiment. SRER = Santa Rita Experimental Range in Arizona; POPL = Poplarville, MS.

## Discussion

The principle objective of this work was to determine whether the exposure of honey bee colonies to low, field-realistic concentrations of clothianidin would have measure effects on colony growth, foraging behavior, thermoregulation and CO_2_ management. Effects have been observed with another neonicotinoid, imidacloprid, at similar exposure rates, in other multi-site experiments [20, 22]. In those studies, honey bee colonies fed 100 ppb imidacloprid in sugar syrup in Arizona, similar to the protocol used here, had lower adult bee populations, brood surface areas and higher within-day temperature variability, compared to colonies in one or more of the other treatment groups, and consumption rates of those colonies were also lower compared to other colonies [22]. In addition, a treatment of 5 ppb imidacloprid affected colonies both at the Arizona site, which was low in alternative forage, as well as at the Mississippi site, rich in alternative forage for much of the year.

In this study discrete (adult bee mass, brood surface area and NEB weights) and continuous (hive weight, internal temperature and CO_2_ concentration) data were collected. Few effects attributable to treatment were observed with respect to continuous data. Variance among treatment groups in terms of hive weight and internal temperature data was mostly explained by the “Experiment” factor, which was a function of time (2017-18 or 2018-19) and place (Arizona or Mississippi). Significant treatment effects were observed only with respect to adult bee mass and the dry weight of newly-emerged bees when two or more experiments were included. These differences correspond to a certain degree with results obtained from other research groups working with the exposure of honey bee colonies to sublethal concentrations of clothianidin [23-27]. The omnibus test for treatment was significant with respect to adult bee mass in the Arizona experiments, but no pairwise comparisons were significant at the α=0.05 level. That NEBs in colonies fed clothianidin 5 ppb were significantly smaller than control NEBs is somewhat unexpected, as it seems to suggest that there were effects at the lowest clothianidin concentration that were not present at the higher concentration. The same effect has been observed at about the same concentrations in another study involving foragers exposed to clothianidin as larvae [3]. Such effects may have been be due to hormesis, defined as a change in the shape of the dose-response curve at low, sublethal concentrations of toxic compounds [33]; effects observed at lower concentrations may be different, or even contrary, to those observed at higher concentrations.

The two Arizona experiments were conducted at the same location, so the kinds of forage would have been the same between those two experiments. However, rainfall was very different between the two years, indicating large differences in the quantity of forage available. The poorer forage opportunities in the 2017-18 season may explain the rapid weight loss in colonies post treatment compared to the following year, and the overall poorer thermoregulation (lower average temperature and greater within-day variability) for colonies from September to December compared to the following year. Similar results were obtained for behavior and thermoregulation of bee colonies given low concentrations of imidacloprid in parallel studies conducted in Arizona and Sydney, Australia [20]. In that study, bee colonies in Sydney, which has considerably higher rainfall than southern Arizona, showed no effect of imidacloprid exposure while those in Arizona did. Additional evidence for reduced alternative forage is provided by the pesticide residue analyses. As with imidacloprid [28], clothianidin was found stable in honey for several months after the end of syrup application. However, while the residues in the original syrup were similar between the two years, the residues from the stored honey were much higher in 2017-18 than in the following year, suggesting less dilution from alternative nectar sources.

Colonies in Mississippi had a strong nectar flow at the beginning of October, as shown by the weight gain among all colonies, and overall better thermoregulation. This may have been due to the more abundant forage in the humid, low-altitude site in Mississippi. Thus, the negative impact of pesticide exposure may have been mitigated by the improved forage in the 2018-19 season compared to the 2017-18 season, and by the overall better forage in Mississippi compared to Arizona.

While continuous hive weight and internal temperature data were not significantly different among treatment groups, continuous CO_2_ concentration data did reveal significant treatment effects in the single season it was deployed. CO_2_ concentration is a function of CO_2_ production and air movement, so one or both of those factors was apparently affected. Like temperature, CO2 concentration in bee hives also are carefully controlled. When [CO_2_], [O_2_] and [N_2_] were manipulated within the hive, only [CO_2_] influenced the fanning behavior of colony members [34]. By controlling CO_2_ concentration in bee hives, bees actively maintain low (15%) [O_2_], causing a reversible hypoxia and reduced metabolic rate among the bees that, researchers have hypothesized, allows them to conserve water and energy, as well as increase activity on short notice [35]. Daily patterns in CO_2_ concentration have been observed in bee hives [36], including peaks of air movement about every 22 seconds [37].

CO_2_ concentration is fundamentally different from measures such as temperature and humidity, which also have ambient (external) counterparts, because ambient CO_2_ concentration is a) very low (on average 409 ppm) compared to internal hive concentrations (>3700 ppm across all treatment groups); and b) varies little (on average about 49 ppm) with respect to time of day compared to the interior of a bee hive (on average >1900 ppm and often >5000 ppm). Ambient temperature and humidity can vary a great deal during the day, and ambient conditions can provide at times higher values than those observed in the hive. That is never the case with the respect to CO_2_ concentration because internal concentrations can never be lower than ambient concentrations. This significant treatment effect suggests further work in understanding the effects of low pesticide concentrations on individual and particularly colony-level behavior. Managing CO_2_ concentration is a vital colony function, and how colonies circulate CO_2_ in the hive likely provides information on colony health.

The importance of landscape in determining colony growth and activity has been observed in several studies. Bee colonies kept in agricultural landscapes were found to have higher growth rates, better thermoregulation, and lower pathogen loads than colonies kept in non-agricultural landscapes [38, 39]. Another study involving commercial colonies in a different set of environments in southern California confirmed those results, and reported better thermoregulation and stronger colonies, in apiaries located in heavy commercial agriculture (Imperial Valley, CA) compared to colonies kept in other landscapes with lower agrochemical exposure [40]. However, in a third set of landscapes, again with commercial colonies, honey bee colonies exposed to commercial agriculture were found to have higher levels of detoxification enzymes and poorer thermoregulation compared to colonies kept on Conservation Reserve Program land [41, 42]. Whether these conflicting results are a result of location-specific factors such as nutritional value of the forage, or reflect unknown factors, remains to be seen. It is hoped that gathering more different kinds of data, on the individual level but particularly on the colony level, such as CO_2_ concentration, might provide further clues in understanding the relationships among bees, landscapes and stressors.

## Acknowledgements

The authors would like to warmly thank M. Heitlinger at the Santa Rita Experimental Range and M. McClaran at the University of Arizona for providing field sites for the work, and R. Scott for access to weather data. In addition, the authors would like to thank M. Alburaki, T. Colin and V. Ricigliano for their helpful suggestions to the manuscript.

---

**S1 File**. Experimental data (XLSX).

**S1 Table**. MANOVA results for the effects of syrup treatment, i.e. clothianidin 20 ppb, clothianidin 5 ppb, and control (blank) across 2 experiments, i.e. Arizona 2017-18 and Arizona 2018-19, and 4 sampling occasions on average adult bee mass (kg) per colony. Hive number was a random factor and pre-treatment adult bee mass was used as a covariate to control for pre-existing differences among colonies.

**S2 Table**. Post hoc contrasts among treatment groups for S1 Table above.

**S3 Table**. MANOVA results for the effects of syrup treatment, i.e. clothianidin 20 ppb, clothianidin 5 ppb, and control (blank) across 3 experiments, i.e. Arizona 2017-18, Arizona 2018-19, and Mississippi 2018-19 and 4 sampling occasions on capped brood surface area (cm^2^) per colony. Hive number was a random factor and pre-treatment adult bee mass was used as a covariate to control for pre-existing differences among colonies.

**S4 Table**. MANOVA results for the effects of syrup treatment, i.e. clothianidin 20 ppb, clothianidin 5 ppb, and control (blank) on Newly Emerged Bee (NEB) dry weights (g) post treatment across 2 experiments, i.e. Arizona 2017-18 and Arizona 2018-19. Ten bees were collected per colony per sampling occasion and the average value per colony was used as the response variable. Hive number was a random factor and pre-treatment NEB dry weight was used as a covariate to control for pre-existing differences among colonies.

**S5 Table**. Post hoc contrasts among treatment groups for S4 Table above.

**S6 Table**. MANOVA results for the effects of syrup treatment, i.e. clothianidin 20 ppb, clothianidin 5 ppb, and control (blank) on Newly Emerged Bee dry weights (g) across 2 post-treatment sampling occasions for the Arizona 2018-19 experiment. Ten bees were collected per colony per sampling occasion and the average value per colony was used as the response variable. Hive number was a random factor and pre-treatment NEB dry weight was used as a covariate to control for pre-existing differences among colonies.

**S7 Table**. Post hoc contrasts among treatment groups for S6 Table above.

**S8 Table**. MANOVA results for the effects of syrup treatment, i.e. clothianidin 20 ppb, clothianidin 5 ppb, and control (blank) across 3 experiments, i.e. Arizona 2017-18, Arizona 2018-19, and Mississippi 2018-19 on hive weight change (g) per colony for days 33-78 after the start of the experiment. Hive number was a random factor and pre-treatment adult bee mass was used as a covariate to control for pre-existing differences.

**S9 Table**. Post hoc contrasts among treatment groups for S8 Table above.

**S10 Table**. MANOVA results for the effects of syrup treatment, i.e. clothianidin 20 ppb, clothianidin 5 ppb, and control (blank) for the experiment conducted in Arizona 2017-18 on hive weight change (g) per colony for days 33-78 after the start of the experiment. Hive number was a random factor and pre-treatment adult bee mass was used as a covariate to control for pre-existing differences.

**S11 Table**. MANOVA results for the effects of syrup treatment, i.e. clothianidin 20 ppb, clothianidin 5 ppb, and control (blank) for the experiment conducted in Arizona 2018-19 on hive weight change (g) per colony for days 33-78 after the start of the experiment. Hive number was a random factor and pre-treatment adult bee mass was used as a covariate to control for pre-existing differences.

**S12 Table**. Post hoc contrasts among treatment groups for S11 Table above.

**S13 Table**. MANOVA results for the effects of syrup treatment, i.e. clothianidin 20 ppb, clothianidin 5 ppb, and control (blank) for the experiment conducted in Mississippi 2018-19 on hive weight change (g) per colony for days 33-78 after the start of the experiment. Hive number was a random factor and pre-treatment adult bee mass was used as a covariate to control for pre-existing differences.

**S14 Table**. MANOVA results for the effects of syrup treatment, i.e. clothianidin 20 ppb, clothianidin 5 ppb, and control (blank) across 3 experiments, i.e. Arizona 2017-18, Arizona 2018-19, and Mississippi 2018-19, on hive internal temperature (°C) for 3 months (1 Sept. – 1 Dec.) after the end of the treatment. Hive number was a random factor and pre-treatment adult bee mass was used as a covariate to control for pre-existing differences.

**S15 Table**. Post hoc contrasts among treatment groups for S14 Table above.

**S16 Table**. MANOVA results for the effects of syrup treatment, i.e. clothianidin 20 ppb, clothianidin 5 ppb, and control (blank) for the experiment conducted in Arizona 2017-18 on hive internal temperature (°C) for 3 months (1 Sept. – 1 Dec.) after the end of the treatment. Hive number was a random factor and pre-treatment adult bee mass was used as a covariate to control for pre-existing differences.

**S17 Table**. MANOVA results for the effects of syrup treatment, i.e. clothianidin 20 ppb, clothianidin 5 ppb, and control (blank) for the experiment conducted in Arizona 2018-19 on hive internal temperature (°C) for 3 months (1 Sept. – 1 Dec.) after the end of the treatment. Hive number was a random factor and pre-treatment adult bee mass was used as a covariate to control for pre-existing differences.

**S18 Table**. MANOVA results for the effects of syrup treatment, i.e. clothianidin 20 ppb, clothianidin 5 ppb, and control (blank) for the experiment conducted in Mississippi 2018-19 on hive internal temperature (°C) for 3 months (1 Sept. – 1 Dec.) after the end of the treatment. Hive number was a random factor and pre-treatment adult bee mass was used as a covariate to control for pre-existing differences.

**S19 Table**. MANOVA results for the effects of syrup treatment, i.e. clothianidin 20 ppb, clothianidin 5 ppb, and control (blank) across 3 experiments, i.e. Arizona 2017-18, Arizona 2018-19, and Mississippi 2018-19, on hive internal temperature amplitudes (°C) from the end of treatment to the end of the annual active season (25 Sept. – 31 Dec.). Hive number was a random factor and pre-treatment adult bee mass was used as a covariate to control for pre-existing differences.

**S20 Table**. Post hoc contrasts among treatment groups for S19 Table above.

**S21 Table**. MANOVA results for the effects of syrup treatment, i.e. clothianidin 20 ppb, clothianidin 5 ppb, and control (blank) for the Arizona 2017-18 experiment on hive internal temperature amplitudes (°C) from the end of treatment to the end of the annual active season (25 Sept. – 31 Dec.). Hive number was a random factor and pre-treatment adult bee mass was used as a covariate to control for pre-existing differences.

**S22 Table**. MANOVA results for the effects of syrup treatment, i.e. clothianidin 20 ppb, clothianidin 5 ppb, and control (blank) for the Arizona 2018-19 experiment on hive internal temperature amplitudes (°C) from the end of treatment to the end of the annual active season (25 Sept. – 31 Dec.). Hive number was a random factor and pre-treatment adult bee mass was used as a covariate to control for pre-existing differences.

**S23 Table**. MANOVA results for the effects of syrup treatment, i.e. clothianidin 20 ppb, clothianidin 5 ppb, and control (blank) for the Mississippi 2018-19 experiment on hive internal temperature amplitudes (°C) from the end of treatment to the end of the annual active season (25 Sept. – 31 Dec.). Hive number was a random factor and pre-treatment adult bee mass was used as a covariate to control for pre-existing differences.

**S24 Table**. MANOVA results for the effects of syrup treatment, i.e. clothianidin 20 ppb, clothianidin 5 ppb, and control (blank) for the Arizona 2018-19 experiment on hive internal CO_2_ concentration (ppm) from 1 Sept to 31 Oct. Hive number was a random factor and pre-treatment adult bee mass was used as a covariate to control for pre-existing differences.

**S25 Table**. Post hoc contrasts among treatment groups for S24 Table above.

**S26 Table**. MANOVA results for the effects of syrup treatment, i.e. clothianidin 20 ppb, clothianidin 5 ppb, and control (blank) on Varroa mite fall post-treatment for the Arizona 2017-18 experiment. Hive number was a random factor and pre-treatment Varroa mite fall was used as a covariate to control for pre-existing differences among colonies.

**S27 Table**. MANOVA results for the effects of syrup treatment, i.e. clothianidin 20 ppb, clothianidin 5 ppb, and control (blank) on Varroa mite fall for two sampling occasions post-treatment for the Arizona 2018-19 experiment. Hive number was a random factor and pre-treatment Varroa mite fall was used as a covariate to control for pre-existing differences among colonies.

## References

1. Mitchell EAD, Mulhauser B, Mulot M, Mutabazi A, Glauser G, Aebi A (2017) A worldwide survey of neonicotinoids in honey. Science 358 (6359):109–111. doi: 10.1126/science.aan3684.

2. Casida JE (2018) Neonicotinoids and other insect nicotinic receptor competitive modulators: Progress and prospects. Annu Rev Entomol 63:125–144. doi: 10.1146/annurev-ento-020117-043042.

3. Morfin N, Goodwin PH, Correa-Benitez A, Guzman-Novoa E (2019) Sublethal exposure to clothianidin during the larval stage causes long-term impairment of hygienic and foraging behaviours of honey bees. Apidologie 50(5):595–605. doi: 10.1007/s13592-019-00672-1.

4. Morfin N, Goodwin PH, Hunt GJ, Guzman-Novoa E (2019) Effects of sublethal doses of clothianidin and/or *V. destructor* on honey bee (*Apis mellifera*) self-grooming behavior and associated gene expression. Sci Rep 9(1) 5196. doi: 10.1038/s41598-019-41365-0.

5. Morfin N, Goodwin PH, Guzman-Novoa E (2020) Interaction of field realistic doses of clothianidin and Varroa destructor parasitism on adult honey bee (*Apis mellifera* L.) health and neural gene expression, and antagonistic effects on differentially expressed genes PLoS ONE 15(2) e0229030. doi: 10.1371/journal.pone.0229030.

6. Tison L, Rößner A, Gerschewski S, Menzel R (2019) The neonicotinoid clothianidin impairs memory processing in honey bees. Ecotoxicol Environ Safety 180:139–145. doi: 10.1016/j.ecoenv.2019.05.007.

7. Abdelkader FB, Kairo G, Bonnet M, Barbouche N, Belzunces LP, Brunet JL (2019) Effects of clothianidin on antioxidant enzyme activities and malondialdehyde level in honey bee drone semen. J Apicult Res 58(5):740–745. doi: 10.1080/00218839.2019.1655182

8. Yao J, Zhu YC, Adamczyk J (2018) Responses of honey bees to lethal and sublethal doses of formulated clothianidin alone and mixtures. J Econ Entomol 111(4):1517–1525. doi: 10.1093/jee/toy140.

9. Henry M, Béguin M, Requier F, Rollin O, Odoux J-F, Aupinel P, et al. (2012) A common pesticide decreases foraging success and survival in honey bees. Science 336:348–350. doi: 10.1126/science.1215039.

10. Samson-Robert O, Labrie G, Chagnon M, Fournier V (2017) Planting of neonicotinoid-coated corn raises honey bee mortality and sets back colony development. PeerJ 8:3670. doi: 10.7717/peerj.3670

11. Tsvetkov N, Samson-Robert O, Sood K, Patel HS, Malena DA, Gajiwala PH, et al. (2017) Chronic exposure to neonicotinoids reduces honey bee health near corn crops Science 356(6345):1395-1397. doi: 10.1126/science.aam7470.

12. Zhu YC, Adamczyk J, Rinderer T, Yao J, Danka R, Luttrell R, et al. (2015) Spray toxicity and risk potential of 42 commonly used formulations of row crop pesticides to adult honey bees (Hymenoptera: Apidae). J Econ Entomol 108(6):2640–2647. doi: 10.1093/jee/tov269.

13. El Hassani AK, Dacher M, Gary V, Lambin M, Gauthier M, Armengaud C (2008) Effects of sublethal doses of acetamiprid and thiamethoxam on the behavior of the honeybee (*Apis mellifera*). Arch Environ Contam Toxicol 54:653–661.

14. Tosi S, Nieh JC, Sgolastra F, Cabbri R, Medrzycki P (2017) Neonicotinoid pesticides and nutritional stress synergistically reduce survival in honey bees. Proc Royal Soc B: Biological Sciences 284:1869. doi: 10.1098/rspb.2017.1711.

15. Coulon M, Schurr F, Martel A-C, Cougoule N, Bégaud A, Mangoni P, et al. (2019) Influence of chronic exposure to thiamethoxam and chronic bee paralysis virus on winter honey bees. PLoS ONE 14(8):e0220703. doi: 10.1371/journal.pone.0220703.

16. Dively GP, Embrey MS, Kamel A, Hawthorne DJ, Pettis JS (2015) Assessment of chronic sublethal effects of imidacloprid on honey bee colony health. PLoS ONE 10(3): e0118748. doi: 10.1371/journal.pone.0118748.

17. Alaux C, Brunet J-L, Dussaubat C, Mondet F, Tchamitchan S, Cousin M, et al. (2010) Interactions between *Nosema* microspores and a neonicotinoid weaken honeybees (*Apis mellifera*). Environ Microbiol 12:774–782.

18. Collison E, Hird H, Cresswell J, Tyler C (2015) Interactive effects of pesticide exposure and pathogen infection on bee health – a critical analysis. Biol Rev 91(4):1006–1019. doi: 10.1111/brv.12206.

19. Pettis JS, vanEngelsdorp D, Johnson J, Dively G (2012) Pesticide exposure in honey bees results in increased levels of the gut pathogen *Nosema*. Naturwissenschaften 99: 153–158.

20. Colin T, Meikle WG, Paten AM, Barron AB (2019) Long-term dynamics of honey bee colonies following exposure to chemical stress. Sci Total Environ 667:660–670. doi: 10.1016/j.scitotenv.2019.04.402.

21. Colin T, Meikle WG, Wu X, Barron AB (2019) Traces of a neonicotinoid induce precocious foraging and reduce foraging performance in honey bees. Environ Sci Technol 53(14): 8252–8261. doi: 10.1021/acs.est.9b02452.

22. Meikle WG, Adamczyk JJ, Weiss M, Gregorc A, Johnson DR, Stewart SD, et al. (2016a) Sublethal effects of imidacloprid on honey bee colony growth and activity at three sites in the U.S. PLOS ONE 11(12): e0168603. doi: 10.1371/journal.pone.0168603.

23. Osterman J, Wintermantel D, Locke B, Jonsson O, Semberg E, Onorati P, et al. (2019) Clothianidin seed-treatment has no detectable negative impact on honeybee colonies and their pathogens Nature Comm 10(1):692. doi: 10.1038/s41467-019-08523-4.

24. Siede R, Meixner MD, Almanza MT, Schöning R, Maus C, Büchler R (2018) A long-term field study on the effects of dietary exposure of clothianidin to varroosis-weakened honey bee colonies Ecotoxicol 27(7):772–783. doi: 10.1007/s10646-018-1937-1.

25. Wood SC, Kozii IV, de Mattos IM, Silva RCM, Klein CD, Dvylyuk I, et al. (2020) Chronic high-dose neonicotinoid exposure decreases overwinter survival of *Apis mellifera* L. Insects 11(1):30. doi: 10.3390/insects11010030.

26. Rolke D, Fuchs S, Grünewald B, Gao Z, Blenau W (2016) Large-scale monitoring of effects of clothianidin-dressed oilseed rape seeds on pollinating insects in Northern Germany: effects on honey bees (*Apis mellifera*) Ecotoxicol 25(9):1648–1665. doi: 10.1007/s10646-016-1725-8.

27. Odemer R, Nilles L, Linder N, Rosenkranz P (2018) Sublethal effects of clothianidin and *Nosema* spp. on the longevity and foraging activity of free flying honey bees. Ecotoxicol 27(5): 527–538. doi: 10.1007/s10646-018-1925-5.

28. Meikle WG, Adamczyk JJ, Weiss M, Gregorc A (2018) Effects of bee density and sublethal imidacloprid exposure on cluster temperatures of caged honey bees. Apidologie 49(5): 581–593. doi: 10.1007/s13592-018-0585-z.

29. Meikle WG, Corby-Harris V, Carroll MJ, Weiss M, Snyder LA, Meador CAD, et al. (2019) Exposure to sublethal concentrations of methoxyfenozide disrupts honey bee colony activity and thermoregulation. PLOS ONE 14(3): e0204635, doi: 10.1371/journal.pone.0204635.

30. Meikle WG, Weiss M (2017a) Monitoring colony-level effects of sublethal pesticide exposure on honey bees. J Vis Exp (129) e56355. doi: 10.3791/56355.

31. Colin T, Bruce J, Meikle WG, Barron AB (2018) The development of honey bee colonies assessed using a new semi-automated brood counting method: CombCount. PLOS ONE 13(10): e0205816. doi: 10.1371/journal.pone.0205816.

32. Meikle WG, Weiss M, Stillwell AR (2016b) Monitoring colony phenology using within-day variability in continuous weight and temperature of honey bee hives. Apidologie 47:1–14. doi: 10.1007/s13592-015-0370-1.

33. Calabrese EJ, Baldwin LA (2003) Hormesis: The dose-response revolution. Annu Rev Pharmacol Toxicol 43:175–97. doi: 10.1146/annurev.pharmtox.43.100901.140223.

34. Seeley TD (1974) Atmospheric carbon dioxide concentration in honey bee (*Apis mellifera*) colonies. J Insect Physiol 20:2301–2305.

35. Van Nerum K, Buelens H (1997) Hypoxia-controlled winter metabolism in honeybees (*Apis mellifera*). Comp Biochem Physiol 117A(4):445–455.

36. Kronenberg F, Heller HC (1982) Colonial thermoregulation in honey bees (*Apis mellifera*). J Comp Physiol B 148:65–76.

37. Southwick EE, Moritz RFA (1987) Social control of air ventilation in colonies of honey bees (*Apis mellifera*), J. Insect Physiol. 33(9), 623–626.

38. Alburaki M, Chen D, Skinner JA, Meikle WG, Tarpy DR, Adamczyk JJ et al. (2018) Honey bee survival and pathogen prevalence: From the perspective of landscape and exposure to pesticides. Insects 9(2), 65. doi: 10.3390/insects9020065.

39. Alburaki M, Steckel SJ, Williams MT, Skinner JA, Kelly H, Lorenz G et al. (2017) Agricultural landscape and pesticide effects on honey bee (Hymenoptera: Apidae) biological traits. J Econ Entomol 110(3): 835–847. doi: 10.1093/jee/tox111.

40. Meikle WG, Weiss M, Beren E (2020) Landscape factors influencing honey bee colony behavior in Southern California commercial apiaries. Sci Rep 10: 5013. doi: 10.1038/s41598-020-61716-6.

41. Meikle WG, Weiss M, Maes PW, Fitz W, Snyder LA, Sheehan T, et al. (2017b) Internal hive temperature as a means of monitoring honey bee colony health in a migratory beekeeping operation before and during winter. Apidologie 48:666–680. doi: 10.1007/s13592-017-0512-8.

42. Ricigliano VA, Mott BM, Maes P, Floyd AS, Fitz W, Copeland DC et al. (2019) Honey bee colony performance and health are enhanced by apiary proximity to US Conservation Reserve Program (CRP) lands. Sci Rep 9: 4894. doi: 10.1038/s41598-019-41281-3.

